# Scalable mapping of myelin and neuron density in the human brain with micrometer resolution

**DOI:** 10.1101/2021.05.13.444005

**Authors:** Shuaibin Chang, Divya Varadarajan, Jiarui Yang, Ichun Anderson Chen, Sreekanth Kura, Caroline Magnain, Jean C. Augustinack, Bruce Fischl, Douglas N. Greve, David A. Boas, Hui Wang

## Abstract

Optical Coherence Tomography (OCT) is an emerging 3D imaging technique that allows quantification of intrinsic optical properties such as scattering coefficient and back-scattering coefficient, and has proved useful in distinguishing delicate microstructures in the human brain. The origins of scattering in brain tissues are contributed by the myelin content, neuron size and density primarily; however, no quantitative relationships between them have been reported, which hampers the use of OCT in fundamental studies of architectonic areas in the human brain and the pathological evaluations of diseases. To date, histology remains the golden standard, which is prone to errors and can only work on a small number of subjects. Here, we demonstrate a novel method that uses serial sectioning OCT to quantitatively measure myelin content and neuron density in the human brain. We found that the scattering coefficient possesses a strong linear relationship with the myelin content across different regions of the human brain, while the neuron density serves as a secondary contribution that only slightly modulates the overall tissue scattering.

## 1. Introduction

The cells, dendrites, and axons in the human brain are structured into cytoarchitectonic and myeloarchitectonic areas, based on cell type, size, density, and the density of myelin sheath surrounding the axons. Those structural components are the substrate for cognitive competencies and the specific locations of neuropathological processes^1–3^. Despite significant advances in imaging technology in the past decades, our understanding of human brain structures at 1-100 μm scale, in which neurons are organized into functional cohorts, is still limited. Quantitative features such as cell and myelin density have only been reported in a small number of subjects and over a small region of the brain^4–9^. Currently, Gallyas Silver stain and Nissl stain are two of the standard histology methods to study myelin content and Neurons in the human brain^10,11^. Despite their ubiquity, complex procedures have to be taken to apply these methods. One needs to cut the brain into tens of micron thickness slices and mount the slice on a glass slide, which induces inevitable tissue damage and distortions. The slices need to be stained and excessive pigment needs to be washed, which is labor intensive and subject to error and variability. Strict control of digitization such as illumination power and camera exposure time are pivotal for downstream quantitative analysis. After imaging individual slices, tremendous efforts are paid to reconstruct the volume, making it challenging for large scale study.

Optical Coherence Tomography (OCT) has been widely used in imaging the brain, which allows volumetric reconstruction of multiple cubic centimeters of tissue with mesoscopic to microscopic resolution. OCT utilizes the backscattered light to acquire tissue microstructural information^12^, which has proven to be useful in revealing cancerous tissue boundaries^13,14^, 3D vascular structures^15–17^, fiber tracts, and individual neurons and laminar structures across the cerebral cortex in rat and human brain^18,19^. OCT also allows quantification of tissue scattering. Fitting the OCT depth profile to a nonlinear model allows the calculation of tissue optical properties such as the attenuation coefficient and the back-scattering coefficient^20,21^. Myelin and neuron scattering have been described as the origins of tissue scattering. Wang et al. 2017 found that the scattering coefficient is higher in white matter and subcortical nuclei regions with highly myelinated fibers, compared to less myelinated grey matter. Srinivasan et al. 2012 found that myelinated fiber tracts are highly scattering while cell bodies have a lower scattering coefficient. Despite these investigations, quantitative correlations between tissue optical properties and these structural components have yet to be investigated.

Here, we report our work on quantifying the relationship between tissue optical properties and myelin content and neuron density in the human brain using automated serial sectioning OCT^22^. We established a computational model of the scattering coefficient with myelin content and cell density as the origins of scattering. By using the ground truth of Gallyas Silver stain and Nissl stain, we showed that the scattering coefficient has a strong linear relationship with the myelin content across different regions of the human brain. We also found that in grey matter, the cell body scattering serves as a secondary contribution to the overall tissue scattering and that the scattering coefficient has a moderate correlation with cell density. Our study provides a novel method for measuring myelin content and neuron density of the human brain tissues in a scalable sample size. The study also has important implications in evaluating brain diseases. As demyelination and neuron loss are two of the pathological hallmarks in neurodegenerative diseases such as Alzheimer’s disease and Chronic Traumatic Encephalopathy (CTE)^23–28^, characterization of the optical property in diseased and normal brains will advance our understanding of pathological evolutions and their impact on complex functions.

## 2. Results

### 2.1 Comparing optical properties with histology

We imaged 5 samples of the human brain with OCT followed by histological staining, which covered anatomical regions of somatosensory cortex, superior frontal gyrus (SupFrontal), middle temporal Brodmann area 21 (BA21), cerebellum, and hippocampus. Gallyas Silver stain and Nissl stain were used as the ground truth measurement of the myelin content and cell bodies in the brain, respectively. For quantitative analysis of myelin content, the Optical Density (OD) of Gallyas Silver stain (Figure 1.a, b, equation (3) in Method) was calculated to represent the density of myelin content in axons assuming the dye concentration is proportional to the myelin density^5,6^. For cell bodies, according to the mie scattering theory (equation (4) in Method), scattering coefficient is proportional to the product of cell density and cross-section area, which we named as cell occupation per area (COPA). COPA was used as the quantitative metric of neuronal scattering (Figure 1.c, d). Optical property maps were estimated from OCT depth profile using a non-linear fitting approach^20^. The scattering coefficient *μ_s_* and the relative back-scattering coefficient 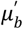 were extracted. Comparing the optical property maps with Gallyas OD and COPA maps allows for inspection of scattering with respect to myelin and neuronal cell body (Figure 1 and Supplemental Figure 1-4).

**Figure 1.**
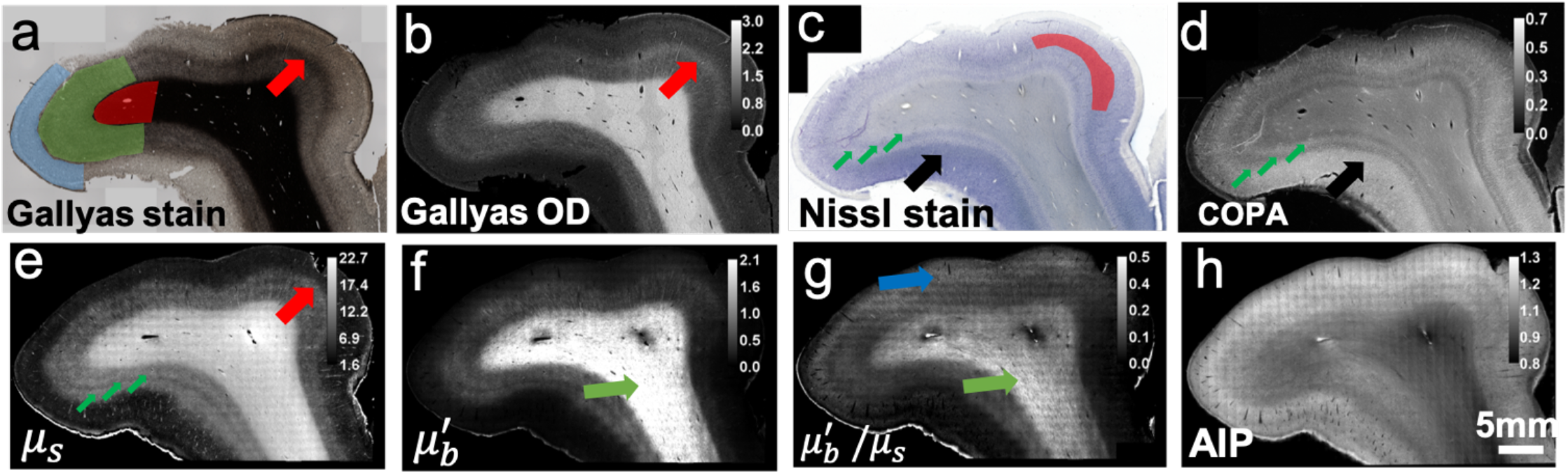
Histology and OCT optical property maps of the human somatosensory cortex. (a) Gallyas Silver stain shows contrast for myelin content. Red region: white matter. Green region: infragranular layers consist of layer IV, V and VI. Blue region: supragranular layers consist of layer I, II and III. Red arrow indicates a thin band of higher myelin content inside layer IV. (b) Optical density (OD) of Gallyas silver stain. The red arrow highlights the myelinated band inside the layer IV. (c,d) Nissl statin and COPA show contrast for cell bodies. The red region indicates the layer II and III with highest COPA value. The big black arrow highlights the high neuron density region. The small green arrows highlight the IV, V, VI layers within the infragranular layer with alternating contrasts. (e-g) Optical properties derived from the OCT images. (e) *μ_s_* map. Small green arrows highlight the alternating contrasts in the infragranular layers similar to that in the COPA map, and the big red arrow indicates the myelinated band seen in Gallyas OD map. (f) *μ*’_*b*_ map. Green arrow highlights the fibers with high intensity, possibly oriented within the imaging plane. (g) Ratio map of *μ*’_*b*_/*μ_s_*. Green arrow highlights the fibers with high intensity, similar to that in the *μ*’_*b*_ map. The Blue arrow highlights the region with high signals in the supragranular layers. (h) AIP image.

Figure 1 shows the histology and optical property maps for somatosensory cortex. Gallyas Silver stain (Figure 1.a) exhibits contrast among the supragranular layers which consist of pyramidal neurons, numerous stellate neurons and sparse axons (indicated by blue region), infragranular layers with large pyramidal neurons and axon bundles that connect to the subcortical structures (green region), and the white matter (red region), which mainly consists of highly myelinated axon bundles and glial cells. The Gallyas OD map (Figure 1.b) demonstrate that the supragranular layers have the lowest OD value, followed by the infragranular layers which have intermediate amount of myelinated axon bundles. The white matter exhibits the highest OD value due to the highly myelinated and densely packed axonal bundles. In addition, smaller features can be seen in the Gallyas OD map as well, such as the thin band of denser myelin content at the upper right region of layer IV (red arrow), possibly due to the high-density fibers in the outer band of Baillarger^29^. The Nissl stain and COPA map (Figure 1.c,d) show contrast for cell bodies. The external granular layer (layer II) and the external pyramidal layer (layer III) exhibit the highest COPA value (red region), especially in the lower left part of the sample (black arrow), probably due to the higher neuron density. The internal granular layer (layer IV), internal pyramidal layer (layer V), and the fusiform layer (layer VI) present alternating contrasts (small green arrows). The white matter generally exhibits a low value of COPA, which is mainly attributed to the glia cells.

The *μ_s_* map (Figure 1.e) strongly resembles the Gallyas OD map. The white matter shows highest *μ_s_* because of the highly scattering myelin sheath. As the sparse axon branches into the cortex, *μ_s_* decreases accordingly. The supragranular layer shows the lowest *μ_s_*, due to the lack of myelin content. In addition, the thin band feature at the upper right region (red arrow) found in Gallyas OD map can also be seen in the *μ_s_* map. Apart from that, the infragranular layers (IV, V, VI) show additional laminar structures (small green arrows) similar to the COPA map but not in Gallyas OD map. Overall, the *μ_s_* map seems to be strongly correlated with myelin content and slightly modulated by the neuron scattering. The *μ*′_*b*_ map (Figure 1.f) offers another feature dimension to aid in discriminating tissue types. It is noticeable that *μ*′_*b*_ varies within the white matter, possibly highlighting fibers oriented within the image plane (green arrow). This is possibly because the fibers oriented within the imaging plane direct more back-scattered photons to the detectors than the fibers oriented through the imaging plane. Consequently, the *μ*′_*b*_ map offers potential information about fiber orientation. The ratio of *μ*′_*b*_/ *μ_s_* map (Figure 1.g) provides another useful feature to distinguish structures in the brain. Similar to the *μ*′_*b*_ map, the ratio of *μ*′_*b*_/*μ_s_* map also highlights the region with fibers oriented within the imaging plane (green arrow). In addition, the ratio of *μ*′_*b*_/*μ_s_* map also highlights region of higher value in the superficial layers indicated by the blue arrow, the cause of such contrast requires further investigation. The average intensity projection (AIP) (Figure 1.h) is a nonlinear function of the *μ_s_* and *μ_b_* maps, which provides an overall view of the tissue structure for reference.

### 2.2 Correlation between scattering coefficient and Gallyas OD

The similarity between the *μ_s_* and Gallyas OD maps indicates that myelin is a crucial factor contributing to the brain tissue scattering. To quantitatively inspect the relationship, we plot *μ_s_* versus Gallyas OD in selective ROIs, which covered all the laminar layers as well as the white matter for the five brain regions. The scatter plots indicate a strong linear relationship between scattering coefficient and Gallyas OD, which is consistent with the mie theory, assuming the same myelin architecture in the tissues^30^, therefore, we fit the data with a linear model and presented the results in Figures 2.a-e.

**Figure 2.**
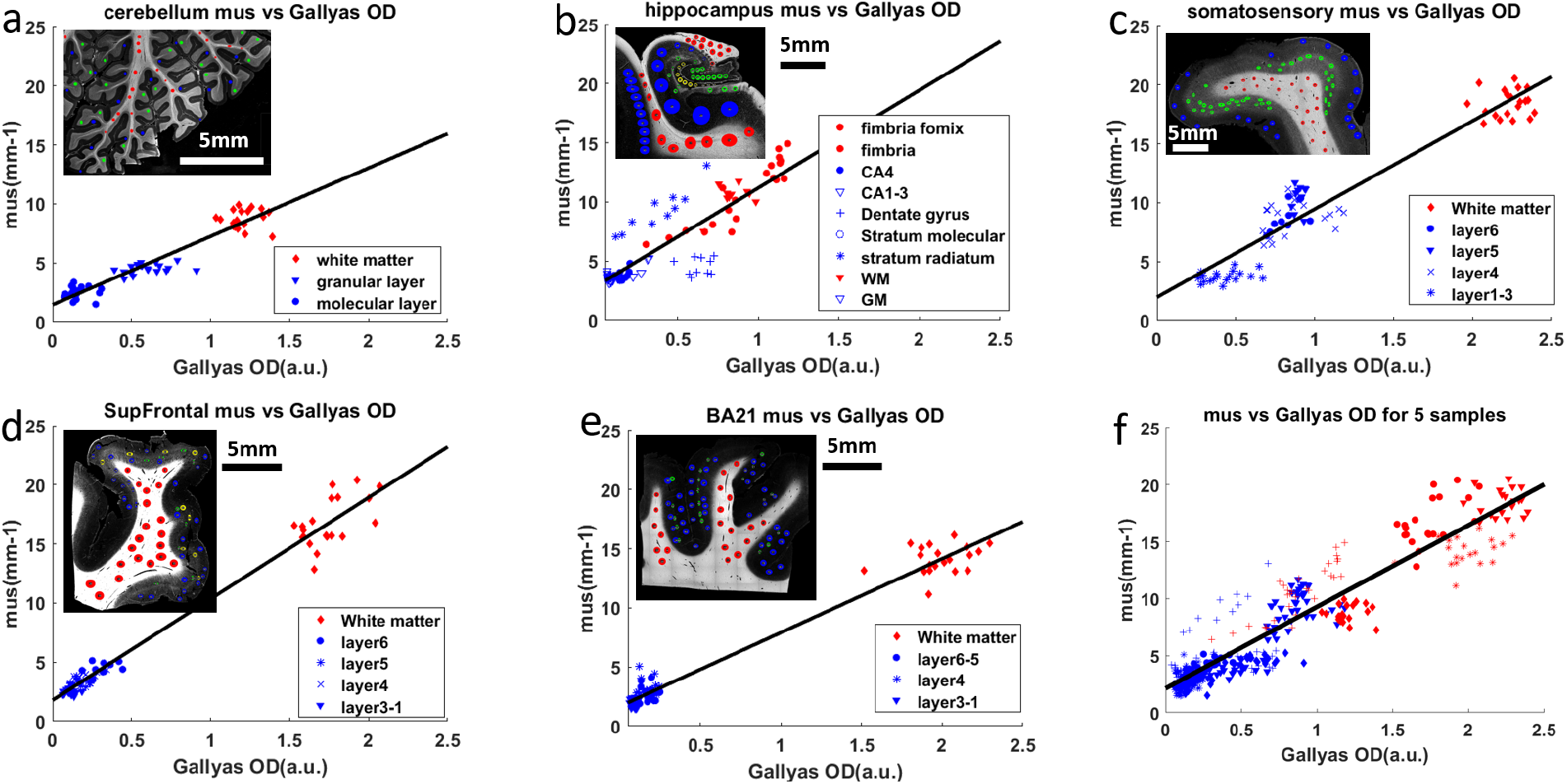
(a-e) Linear regression of *μ_s_* and Gallyas OD for 5 different regions in human brain samples. Red dots: white matter data points. Blue dots: grey matter data points. The inset figure shows the Gallyas OD map of the corresponding sample. The Red circles in the inset figure represent the ROIs in the white matter. The blue, green and yellow circles represent the ROIs in different layers of the grey matter, for example, the green ROIs in (c) represent infragranular layers and the blue ROIs represent the supragranular layers. (a) cerebellum. (b) hippocampus. (c) somatosensory cortex. (d) Superior frontal cortex (SupFrontal). (e) middle temporal Brodmann area 21(BA21). (f) linear regression of all data points from 5 samples. The six panels have the same range on the X and Y axes for easier comparison.

Remarkably, the five samples share similar linear relationships as indicated by the slope parameter (*k*_1_) of the fitting results, although individual samples have distinct patterns of *μ_s_* and Gallyas OD distributions. In the cerebellum (Figure 2.a), data points from the white matter, granular layer and molecular layer form three discrete clusters. However, they all follow a shared linear function (*k*_1_ = 5.79) in which higher Gallyas OD is associated with higher *μ_s_*. Similar patterns are observed in the somatosensory cortex (Figure 2.c), where the six cortical layers and the white matter group into three clusters and share a co-linear relationship (*k*_1_ = 7.48). Supragranular layers (layer I, II and III) form a cluster with the lowest Gallyas OD and *μ_s_*, infragranular layers (layer IV, V and VI) form a cluster with intermediate values, and the white matter cluster exhibits the highest values. The other two cortical regions of SupFrontal and BA21 (Figures 2.d-e) only present two discrete clusters. The supragranular layer and infragranular layer in both regions fall into a single cluster with low Gallyas OD and *μ_s_*. The white matter tracts show high values close to those of somatosensory cortex. Interestingly, the SupFrontal displays a within-cluster trend in both grey matter and white matter, suggesting a myelin gradient across cortical layers. Regardless, the slope parameters (*k*_1_ = 8.57 for SupFrontal and *k*_1_ = 6.24 for BA21) demonstrate a similar relationship as those revealed in the cerebellum and the somatosensory cortex. In the hippocampus (Figure 2.b), due to complex anatomical structures, data points from different layers form a continuous distribution. For example, the fimbria, white matter and fornix show gradually increasing Gallyas OD and *μ_s_* values, while having large overlaps with the CA4 and dentate gyrus. Despite the different distribution pattern from other tissues, the fitting result is comparable with a slope parameter *k*_1_ = 8.26.

An extraordinary linear relationship between Gallyas OD and *μ_s_* is revealed in all the samples (P<10^-10^) with high correlation coefficients (PCC>0.85 for all brain regions, see Figure 3.b). The slope between Gallyas OD and *μ_s_* falls in a narrow range of 5.8 to 8.6. Slope variations are possibly due to different staining backgrounds, as well as other scattering factors such as neuronal cell bodies. In Figure 2.f we combined all the data from the 5 samples and fit a single linear function, which reveals an average slope of 7.2 (correlation coefficient = 0.936, P<10^-60^). These results provide evidence that the linear relationship between optical scattering and myelin content holds in different regions of the brain. Therefore, scattering coefficient may potentially be used as an objective measurement of the myelin content in the human brain.

**Figure 3.**
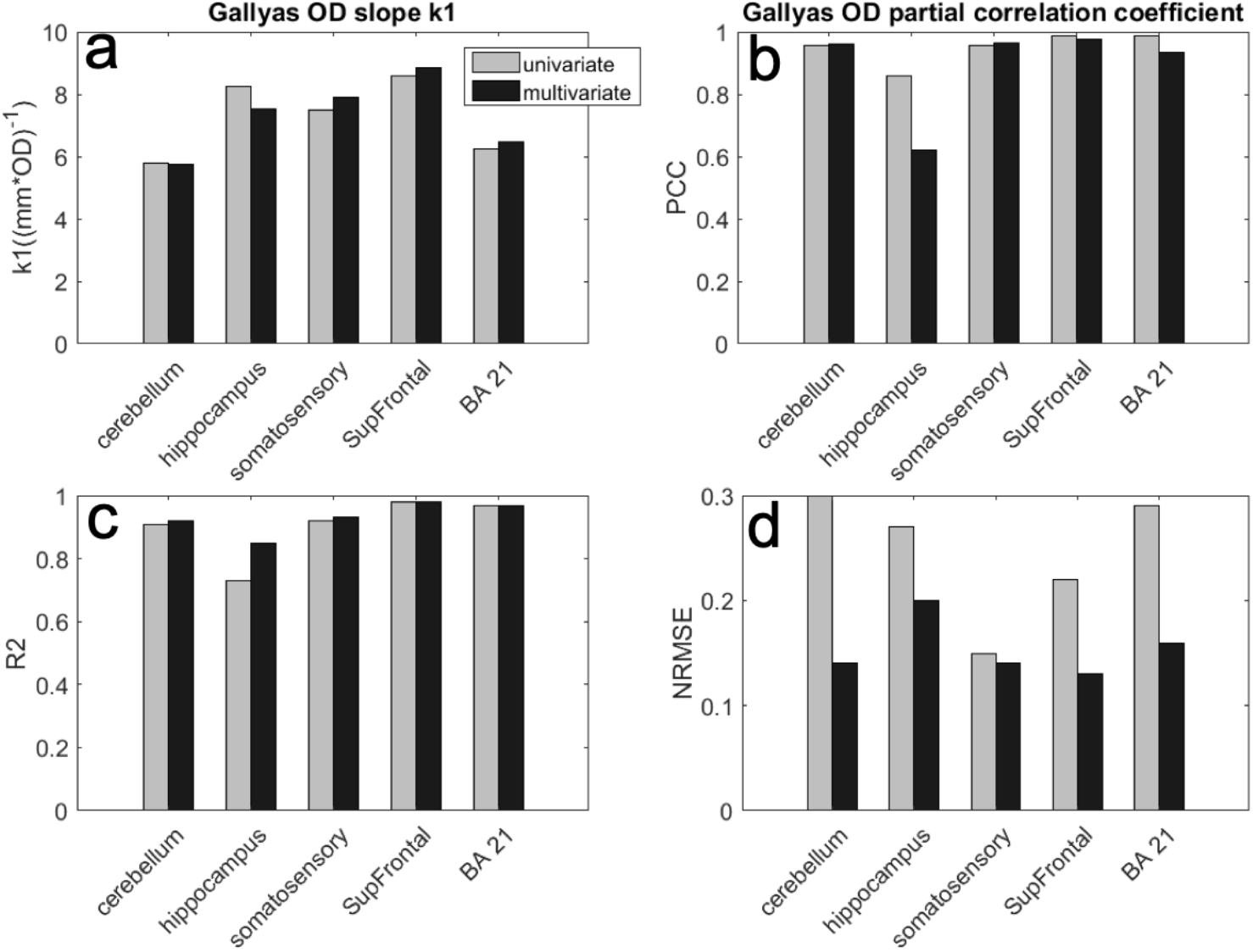
Multivariate regression of *μ_s_* vs Gallyas OD and COPA (grey bars), compared against univariate regression where only Gallyas OD is considered (black bars). (a) Gallyas OD slope *k*_1_ of 5 brain regions resulted from univariate regression and multivariate regression. From left to right: cerebellum, hippocampus, somatosensory, SupFrontal and BA21. (b) Correlation coefficient with Gallyas OD in univariate and partial correlation coefficient in multivariate regression. (c) R^2^ of Pearson’s correlation in univariate and multivariate regressions. (d) Normalized root mean square error (NRMSE) of univariate and multivariate regressions.

### 2.3 Correlation between scattering coefficient and joint Gallyas OD and COPA

The results of the linear model in session 2.2 suggest that myelin is a strong predictor of scattering coefficient in both grey and white matter of the human brain. In the cortex, neuronal cell body is another factor that should be taken into consideration when interpreting the scattering coefficient of the brain tissue. In this section, we first re-evaluate the relationship of *μ_s_* and Gallyas OD by including the factor of COPA. Next, we quantify the correlation between *μ_s_* and COPA to examine the cell body contribution in scattering. For such purpose, we built a multivariate, generalized linear model to include both Gallyas OD and COPA as predictors (equation (5–8) in Method). In the grey matter, we included both Gallyas OD and COPA as contributing factors, whereas in white matter, we only considered Gallyas OD since contribution from neuronal cell bodies was neglectable.

Figure 3.a compares the slopes for Gallyas OD (*k*_1_) in univariate regression where only Gallyas OD was considered, and multivariate regression where COPA was included as well. We found that *k*_1_ values were highly comparable between these two models with differences less than 10% in all brain regions Besides, the partial correlation coefficient (PCC) of Gallyas OD in the multivariate regression (Figure 2.d) was found higher than 0.9 for four out of five samples (P<10^-10^ for all), with a slightly lower PCC of 0.6 in the hippocampus. Compared to the univariate regression, the partial correlation coefficient in the multivariate regression has only minor difference in most samples. The consistency of *k*_1_ in the two analyses reinforces that the linear relationship between *μ_s_* and Gallyas OD largely preserves even when cell body scattering is taken into account. The high PCC in multivariate model consolidate the finding in session 2.2 that *μ_s_* is strongly correlated with myelin content across brain regions. In addition, when evaluating the R^2^ of Pearson’s correlation and Normalized Root Mean Square Error (NRMSE, Figure 3.c,d) for the regression, we found that adding COPA into the model only results in small improvements, which indicates that myelin content is a dominant factor to scattering coefficient.

Figure 4 elaborates the correlation between *μ_s_* and COPA in the multivariate regression model. The slope for COPA (*k*_2_) exhibits large variations across the 5 different samples (Figure 4a). In somatosensory cortex, Supfrontal and BA21, a positive *k*_2_ was found, while in cerebellum and hippocampus, *k*_2_ was negative. Two-sided t-test reveals a significant positive correlation only in the somatosensory region (P<0.01, PCC=0.45). The correlations in SupFrontal and BA21 are moderate (Figure 4b), but further statistical test fails to find a significance (P=0.11 for SupFrontal and P=0.35 for BA21). These results suggested that the neuronal scattering is a small contribution to the overall scattering coefficient and the effect varies across the brain. The negative correlation in the cerebellum and hippocampus was counterintuitive. However, it should be noted that the size of neurons in densely packed layers such as the granular cells in the cerebellum and the dentate gyrus in the hippocampus is much smaller than that of the other layers, which leads to a reduced scattering coefficient. The intercept in the multivariate regression exhibits large variation as well (Figure 4c). As the intercept in the regression encompasses the unmodeled components to the tissue scattering, such as the extracellular matrix, the small intercepts in hippocampus, somatosensory, and BA21 indicate a negligible contribution from these remaining components. However, in cerebellum and SupFrontal, we found a significant intercept (P<0.001), indicating substantial scattering components remained. Overall, the fitting results for COPA in the model are coherent with findings in session 2.2 and 2.3 that *μ_s_* is dominant by the myelin factor. Neuronal cell body, however, only plays a secondary contribution to tissue scattering in the human brain.

**Figure 4.**
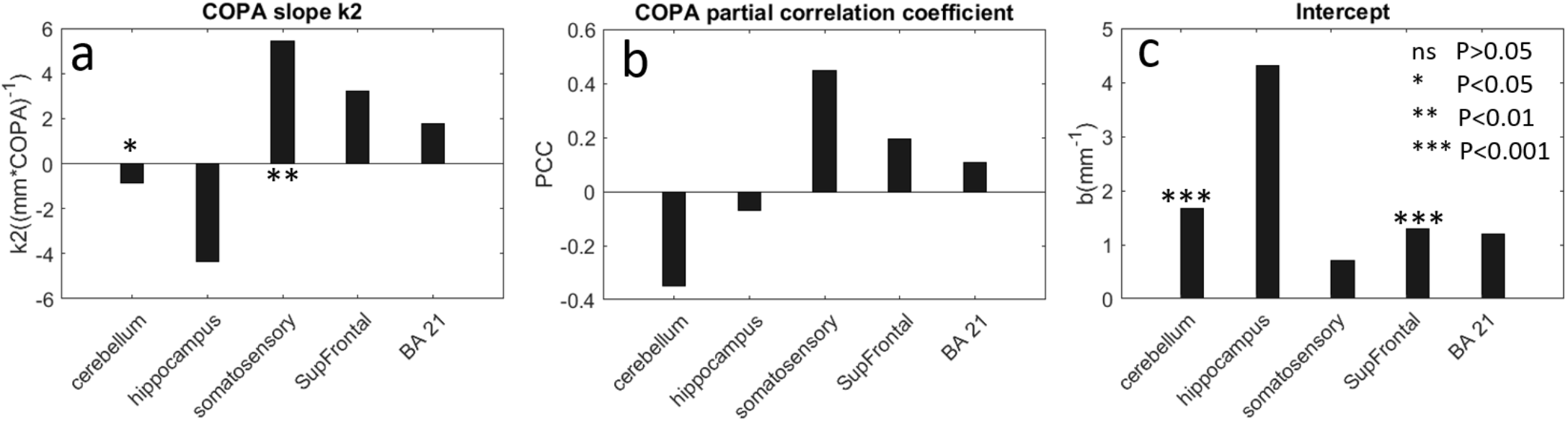
Evaluation of *μ_s_* with COPA and remaining factors in multivariate regression. (a) COPA slope *k*_2_ in the five brain regions. Regions with stars indicate significant *k*_2_. (b) Partial correlation coefficient of COPA with respect to *μ_s_*. (c) Intercept b of the multivariate regression in the five brain regions.

### 2.4 Correlation of *μ*′_*b*_ and *μ*′_*b*_/*μ_s_* with COPA and Gallyas OD

We analyzed the relationships of *μ*′_*b*_ and *μ*′_*b*_/*μ_s_* with respect to the COPA and Gallyas OD, respectively. For cerebellum, somatosensory cortex, BA21 and SupFrontal, there is a strong linear relationship between *μ*′_*b*_ and Gallyas OD (Supplemental Figure 5,8), which is likely a result of the strong dependency between *μ_s_* and *μ*′_*b*_. However, there is no clear relationship between the ratio map of *μ*′_*b*_/*μ_s_* and Gallyas OD or COPA (Supplemental Figure 6,7).

### 2.5 Scattering coefficient differentiates neocortex from allocortex

As shown in Figure 2, *μ_s_* and Gallyas OD exhibit distinctive cross-layer patterns among the 5 samples. As both metrics are strong predictors of myelin content, we examined the mean *μ_s_* and Gallyas OD in the white matter (red ROIs and red dots on scatter plots of Figure 2) for all the samples. We found that *μ_s_* distribution exhibits a similar pattern as in Gallyas OD (Figure 5a-b). Hippocampus and cerebellum have significantly lower Gallyas OD and *μ_s_* compared to the other three samples in the cortex, indicating a lower myelin content in the two regions of the brain. Anatomically, cerebellum and hippocampus belong to the allocortex while somatosensory, SupFrontal and BA21 belong to the neocortex. The two types of cortices possess different developmental trajectories of myelination (Miller et al., 2012). The mean COPA in the grey matter (blue dots on scatter plots of Figure 2) of the allocortex is significantly higher than that of the neocortex (Figure 5.c). Our results suggest that in addition to histology, the optical property obtained by OCT serves as a viable tool to differentiate the neocortex from the allocortex, with a distinction resulting from underlying myelin content.

**Figure 5.**
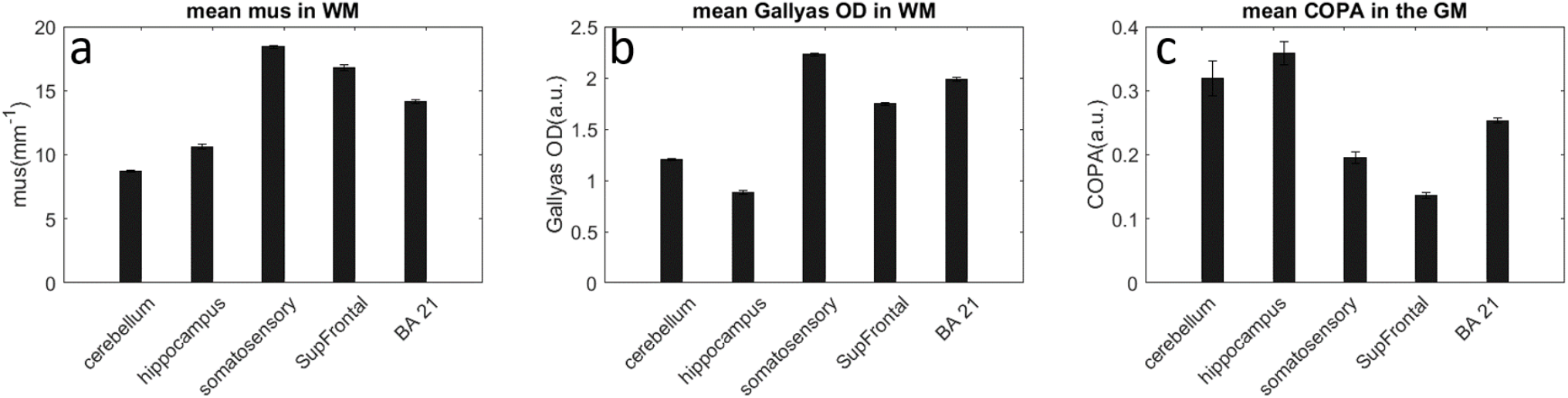
(a) average *μ_s_*, (b) average Gallyas OD in white matter and (c) average COPA in grey matter differentiating brain regions. The error bars represent the standard error from the ROIs.

## 3. Discussion and Conclusions

Previous studies have demonstrated the ability of OCT to differentiate cortical laminar structures and to identify fiber tracts and subcortical nuclei^20,31^. Here, by using serial sectioning OCT, we have quantitatively investigated the contribution of structural components to the optical property in human brain samples and established a model of the scattering coefficient with regard to myelin content and neuron density. We have found that the scattering coefficient is strongly correlated with the myelin content (P < 10^-10^, PCC > 0.85) and that linear relationship is retained across different regions of the brain. The domination of myelin content in tissue scattering is reasonable considering the high index of refraction of myelin (n=1.47) with respect to the surrounding aqueous environment (n=1.35) (Kwon et al., 2017). The results from our study suggest that the optical property can be used a robust predictor for myelin content of the human brain. This strong correlation between scattering coefficient and myelin content has important implications in neurodegenerative diseases. It has been shown that the breakdown of myelin sheath is an indication of pathological abnormality and can result from several neurodegenerative diseases such as multiple sclerosis^32–34^, Alzheimer’s disease^25,35^ and chronic traumatic encephalopathy^23,24^. The quantitative measurement of myelin content could potentially be useful in characterizing the degree of demyelination in pathological brain samples. In addition, we have shown that scattering coefficient enabled the differentiation of various brain regions, such as neocortex and allocortex that have distinct myelination trajectories in development^36^. Previous studies have revealed that the degree of myelination and the order of maturation in the brain is associated with the vulnerability to psychiatric disorders^37–39^. Systematic characterization of myelination in the brain may provide a new avenue to map out the regional vulnerability to a range of brain disorders.

Neuronal cell body turns out to be a secondary contribution to the overall scattering, and the correlation varies across different brain regions. In somatosensory cortex we found a significantly positive correlation (P<0.01), indicating a strong laminar structure with differed neuron density and size, while in other brain regions we observed negative or moderately positive correlations. The lack of major contribution made by cell bodies to scattering in brain tissues have been reported in previous studies. Kalashnikov et al. found light scattering from the neuron body contributes less than 10% of the observed backscattering signal when using cultured Hela neurons. Besides, Magnain et al. found out that myelin density and fiber orientation could disrupt the identification of neuron cell bodies by using an optical coherence microscopy (OCM). As a result, the weak correlation revealed by the current linear model is not unexpected. Indeed, most of the OCT studies on brain cancer samples have reported difficulties in differentiating cancer of various stages from normal grey matter merely by scattering coefficient^42,43^. Our studies might provide an explanation for those challenges because cell body contribution is only a minor factor for light scattering in the brain. Further improvement of our scattering model may increase the sensitivity to neuron scattering. In our model, we assumed the same k1 and intercept between the grey matter and white matter (Figure 2 and Supplementary figure 9). However, as the refractive index of the extracellular space in grey and white matter could be different, allowing parameter tuning might result in a better fitting. When formulating the relationship between scattering coefficient and COPA, we assumed the phase function Qs to be constant for all neuron bodies (equation. 4, Method), which bears a drawback if there’s a large variation in the neuron size. Considering the dependency between phase function and scatterer size, a nonlinear model might improve the performance for correlating scattering coefficient with neuronal cell bodies.

Compared to histological methods, serial sectioning OCT offers a new window for quantifying myeloarchitecture and neuroarchitecture in human brain. Histological stains have been standard methods for studying myelin content and cell density. However, the outcome of histology heavily depends on the concentration of the contrast agent, the pH and temperature, and bleaching procedure for undesired pigment, which may vary across studies. Histology also requires a series of manual processes that are prone to human errors. Consequently, variations from slice to slice are inevitable^44^. In addition, sample must be cut into thin slices before being stained and imaged, which introduces tissue damage and distortions that are challenging to correct during 3D reconstruction. Serial sectioning OCT, on the contrary, uses intrinsic optical properties of tissue that does not depend on external contrast agents. The scattering coefficient is insensitive to system setup, incident power, and acquisition parameters. In volumetric imaging, the images are acquired on blockface prior to slicing, avoiding the vast majority of distortions incurred by tissue cutting and mounting. As a result, serial sectioning OCT generates images with consistent qualities across slices and samples. The technique, being automated and robust, is favorable to expand to larger sample size. Thus, serial sectioning OCT provides an attractive solution for quantifying volumetric myelin content and neuron cells in the human brain.

A few future directions in this field can be pointed out. First, another optical property called birefringence may directly relate to myelin content in the brain^45^. With polarization-sensitive OCT, we can measure the birefringence of myelinated fibers in addition to the scattering coefficient^46^. Inclusion of both parameters will enable a model for predicting and synthesizing myelin content in the human brain. Second, the local index of refraction may serve as another optical property to quantify neuronal characteristics^19^. Besides, high-resolution OCM has proven to be able to visualize neurons in brain tissue. Magnain et al. used OCM to identify neurons that were validated by co-registered Nissl stain images in human entorhinal cortex. In addition to OCT, other imaging techniques such as two photon microscopy (2PM) could be useful to quantify neuron density as well. With elongated depth of profile, 2PM is able to cover a volume of tissue with high volume rate and generates an average intensity projection (AIP) of autofluorescence signals^47–50^. Hence, one of the future directions is to use multimodal techniques to obtain accurate measurement of cell and myelin content in scalable human brain samples.

Several empirical considerations ought to be clarified in this study. First, in the effort of correlating to histology, we thoroughly examined the quality of histological images and used the slices that have consistent staining intensity as ground truth. Yet there might be minor variations of staining that were not normalized among different samples, which could be one of the reasons for the variations of the slope and b values observed in the fitting results. Second, we formulated COPA based on the assumption that neuron bodies do not overlap on Nissl images, which may result in an underestimation in regions with high neuron populations. As an alternative, we also calculated the OD of Nissl stain and correlated it with the optical properties. The results were not significantly different from using COPA. Third, to obtain COPA we used an empirical thresholding method to segment the neuron body while excluding the smaller glia cells, the accuracy of which may depend on brain regions and staining quality. In the future, a deep learning based classifier may improve the segmentation^9^

In conclusion, we have demonstrated the use of optical scattering obtained by serial sectioning OCT to measure myelin content and neuron density of human brain tissues. The scattering coefficient has a strong linear relationship with the myelin content across different brain regions, which promotes a robust label-free measurement for myelin with substantially reduced labor efforts comparing to traditional histology. The scattering coefficient was also moderately modulated by the neuronal cell bodies, the precise measurement of which requires complementary quantification or imaging techniques. Our approach has great potentials for enabling large-scale investigation of myeloarchitecture in various brain regions as well as studies of neurodegenerative processes in pathological brain samples.

## Method

### Sample

Two human brains (mean age 53.5 ± 12.0 y.o., 1 male and 1 female) were obtained from the Massachusetts General Hospital Autopsy Suite. The brains were neurologically normal without a previous diagnosis of neurological deficits. The tissues were fixed by immersion in 10% formalin for at least two months^51^. The post-mortem interval did not exceed 24 h. The samples in the study consisted of five 5 anatomical regions from the two2 human brains, including the cerebellum, hippocampus, somatosensory cortex, superior frontal cortex, and middle temporal area 21. Each sample was embedded in 4% melted oxidized agarose and cross-linked with a borohydride-borate solution for block-face imaging^52^.

### Serial Sectioning OCT system

The serial sectioning OCT system was previously described^22^. Briefly, the system consists of a spectral-domain OCT system to measure the optical properties of the sample as well as a customized vibratome for tissue slicing. The light source was a broadband super-luminescent diode with a center wavelength of 1300 nm and full width half maximum bandwidth of 170 nm, yielding an axial resolution of 3.5 μm in tissue. The spectrometer consisted of a 1024-pixel InGaAs line scan camera operating at an A-line rate of 47kHz. The total imaging depth was estimated to be 1.5 mm. The sample arm used a 10 × water immersion objective (Zeiss, N-Achroplan), yielding a lateral resolution of 3.5μm. The volumetric imaging covered a field of view (FOV) of 1.5 × 1.5 × 1.5 mm^3^. The voxel size, which was defined by the stepping size of galvo mirror scanning laterally and obtained by the total imaging depth divided by the number of pixels axially, was 2.9 μm isotropic. The sensitivity of the system was 105 dB. Consecutive image tiles were obtained to cover the entire area of the sample and a 50% overlap was used between adjacent tiles. A customized vibratome was mounted adjacent to the OCT to cut off a 50μm thick slice of the tissue upon completion of the full area scan. The slices were retrieved for histological staining. It is noted that the comparison of OCT and histology images in this study was not conducted on the same slices but slices nearby, as OCT slices have been preserved for other uses.

### Quantification of optical properties

Optical property maps from the OCT data of the five human brain samples are estimated using the non-linear model based depth profile fitting approach described in^20,53^. Specifically, we used the following equation to fit the depth profile of the OCT signal, from which we extracted the scattering coefficient *μ_s_* and the relative back-scattering coefficient 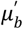. Note that 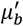 is a relative variable, depending on factors of the OCT system including the incident power and the spectrometer efficiency, and is proportional to the intrinsic back-scattering coefficient *μ_b_*:

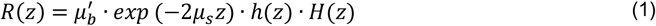

where R(z) is the OCT reflectance signal versus depth, H(z) is the sensitivity roll-off function^20^, h(z) is the axial point spread function of the microscope system, which depends on the focus depth *Z_f_* and the effective Rayleigh range *Z_RS_* of the objective:

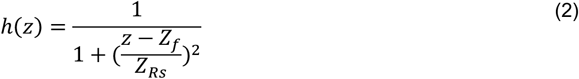

In general, the fitting process tries to find the optimal value for four parameters: 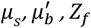 and *Z_Rs_* for each A-line. Without any fitting constraints there is a strong inter-dependency between these parameters^20^. To reduce this inter-dependency and to make the fitting more stable and robust, we spatially parameterized the *Z_f_* and *Z_Rs_* following the procedure described in Yang et al., 2020. The *Z_f_* and *Z_Rs_* values were pre-calibrated using an Intralipid phantom with a comparable scattering coefficient as the tissue sample. As a result, we have reduced the number of unknowns in our fitting model to two: *μ_s_* and 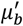. To reduce the noise and errors in estimating these optical properties, the OCT A-lines were averaged over a 30×30 μm^2^ area before fitting. A ratio map of 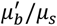 was obtained afterward.

In reconstruction, we used ImageJ to stitch the Average Intensity Projection (AIP) of each OCT image tile to generate the XY coordinates for overlapping tiles. The tiles were then blended using a customized Matlab code to remove artifacts caused by intensity non-uniformity across the field of view. The down-sampled (30×30 μm^2^, same as the volumetric averaging in fitting) stitching coordinates were used to stitch the *μ_s_* and 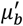 results from previous fitting steps.

### Histology and digital image processing

Selected slices from the serial block-face scanning were processed with Nissl stain^54,55^ for neuron body identification and Gallyas stain^10,36^ for characterizing myelin content. The stained slices were digitized by an 80i Nikon Microscope (Micro-video Instruments, Avon, Massachusetts) with a 4x objective. The pixel size was 1.9 μm.

The Gallyas stain images were normalized in each channel and converted to mean Optical Density (OD)^44^ to directly represent the myelin content:

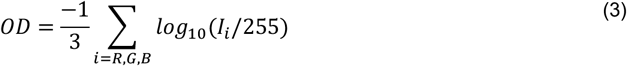

where *I* is the RGB vector of the histology image with *I_i_* representing the intensity of the red, green, or blue channel, respectively. Each RGB channel was normalized separately before converting to OD in log scale. Then the average of three channels was used as the mean OD that represents the myelin content.

For Nissl stain, the percent of the local area occupied by neuron bodies (COPA) was computed in two steps: first, the neuron bodies were segmented by converting the original image to a binary image in ImageJ using the binary function. The binary image was then downsampled by 10×10 pixels to get the localized neuron body occupation percentage. This quantity is proportional to the cross-sectional area of the neurons locally, which, according to Mie theory^56^, is related to the scattering coefficient. Specifically, the scattering coefficient of a sphere is given by

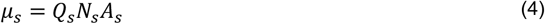

Where *Q_s_* is the phase function depending on the scatterer size, wavelength of the light source, and scattering direction; *N_s_* is the number density of the scatterer; and *A_s_* is the cross-sectional area of the scatterer^56^. Assuming that the phase function *Q_s_* is constant for all neuron bodies, the scattering coefficient becomes proportional to the product of cellular density and cellular cross-sectional area, which we define as the cellular occupation per area (COPA). Here we assume that the neuron bodies do not overlap in the 2D projection histology images. The validity of such an assumption is explained in the limitation part of the discussion.

### Generalized linear model

We examined the relationship between the optical properties and the Gallyas OD and COPA to reveal the sources of tissue scattering. OCT block-face and the histology images were acquired on slices from the same brain region that were a few millimeters apart. We manually drew 60 to 90 Region of Interests (ROIs) on corresponding slices from both modalities for linear regression analysis. The area of ROIs ranges from 200 pixels to 500 pixels. In larger areas with more uniform intensity, such as the grey matter in hippocampus, the area of ROIs were expanded to a few thousand pixels to get more precise measurement. The total number of ROIs depends on the number of distinct cortical layers or subdivisions. On average we drew 20 evenly distributed ROIs for each layer in the *μ_s_*, Gallyas OD and COPA maps, respectively.

We used the generalized linear model (GLM) to study the relationship between tissue scattering, myelin content and neuron scattering. The GLM relationship can be mathematically described as,

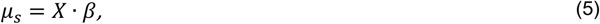

Where the matrix *X* contains the measured Gallyas OD and COPA values and *β* contains the linear coefficients that will be estimated:

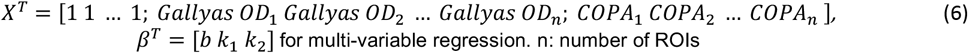

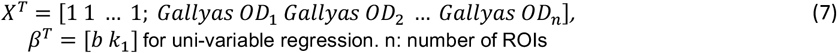

where *k*_1_ is the slope corresponding to the myelin contribution to the tissue scattering, *k*_2_, is the slope corresponding to the neuronal contribution to the tissue scattering, and b is the contribution from other components, such as scattering from the extracellular matrix. The coefficients were calculated by a pseudoinverse solution as,

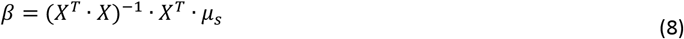

We used *R*^2^ of Pearson’s correlation and Normalized Root Mean Square Error (NRMSE) for examining the goodness of the fitting. Partial correlation coefficient (PCC), two sided t-tests and multiple comparisons were conducted to reveal the significance of the contributions from Gallyas OD and COPA, respectively.

It should be noted that we set COPA in the white matter of all tissue types to be 0 before regression, as there are only glia cells and a few interstitial neurons^57^ in the white matter, which won’t contribute to scattering significantly. Leaving it unchanged, however, will bias the regression. In the multivariable regression, we also assumed that *k*_1_ and b to be the same in grey and white matter, as we assume that the contribution from myelin and other extracellular components behave similarly, and thus use the same coefficients *k*_1_ and b to describe them jointly.

Lastly, we also investigated the origins of contrast in the 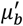 map and ratio of 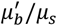. Four linear models were used, namely, fitting 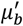 as a linear function of Gallyas OD and COPA, respectively, and fitting the ratio of 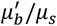 as a linear function of Gallyas OD and COPA, respectively. Due to the lack of strong correlation of 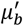 or ratio of 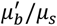 with respect to Gallyas OD and COPA (see session 2.4 and supplementary figures), no multivariate investigation was further performed.

## Data Availability

The datasets generated and/or analyzed during the current study are available from the corresponding author on reasonable request.

## Acknowledgements

Support for this research was provided in part by the BRAIN Initiative Cell Census Network grant U01MH117023, the National Institute for Biomedical Imaging and Bioengineering (P41EB015896, 1R01EB02328, 1R01EB006758, R21EB018907, R01EB019956, P41EB030006), the National Institute on Aging (1R56AG064027, 1R01AG064027, 5R01AG008122, R01AG016495), the National Institute of Mental Health (R01 MH123195, R01 MH121885), the National Institute for Neurological Disorders and Stroke (R01NS0525851, R21NS072652, R01NS070963, R01NS083534, 5U01NS086625, 5U24NS10059103, R01NS105820), a NIH career development award R00EB023993 to HW and grant number 2019-189101 from the Chan Zuckerberg Initiative DAF, an advise fund of the Silicon Valley Community for CM, and was made possible by the resources provided by Shared Instrumentation Grants 1S10RR023401, 1S10RR019307, and 1S10RR023043. Additional support was provided by the NIH Blueprint for Neuroscience Research (5U01-MH093765), part of the multi-institutional Human Connectome Project.

## Author contributions

S.C., D.A.B., and H.W. conceived and designed the research; C.M., J.A. and H.W. performed data acquisition; S.C. analyzed the data and constructed the mathematical models with input from D.V. and D.G.; D.V., J.Y, A.C., S.K., C.M., J.A., B.F., D.G., D.A.B. and H.W. contributed to data analysis and interpretation; S.C., D.V., and H.W. contributed to manuscript preparation.

