## Supplemental Figures for "Scalable mapping of myelin and neuron density in the human brain with micrometer resolution"

**Supplementary figures**

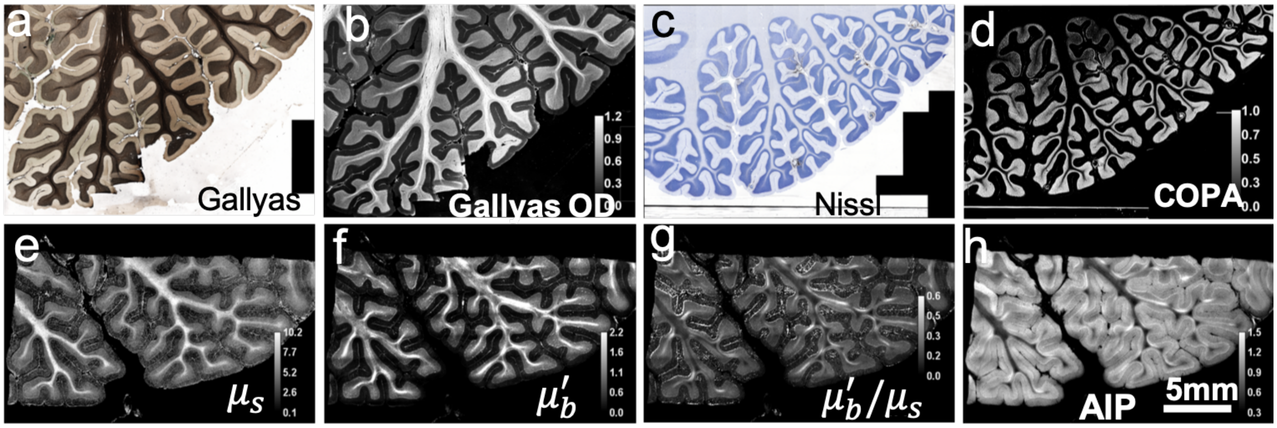


Supplementary Figure 1. Histology and optical properties of cerebellum. (a) Gallyas Silver stain. (b) Optical density of Gallyas silver stain reveals the myelin content. (C) Nissl stain. (d) COPA. (e-g) Optical properties derived from the OCT images. (e) $\mu_s$ map. (f) $\mu'_b$ map. (g) ratio map of $\mu'_b / \mu_s$. (h) AIP image

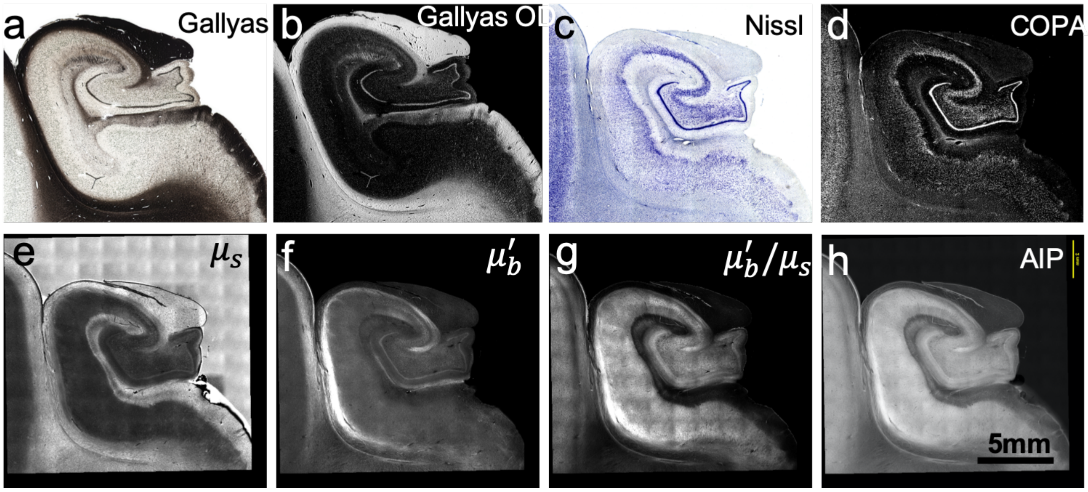


Supplementary Figure 2. Histology and optical properties of Hippocampus. (a) Gallyas Silver stain. (b) Optical density of Gallyas silver stain reveals the myelin content. (C) Nissl stain. (d) COPA. (e-g) Optical properties derived from the OCT images. (e) $\mu_s$ map. (f) $\mu'_b$ map. (g) ratio map of $\mu'_b / \mu_s$. (h) AIP image

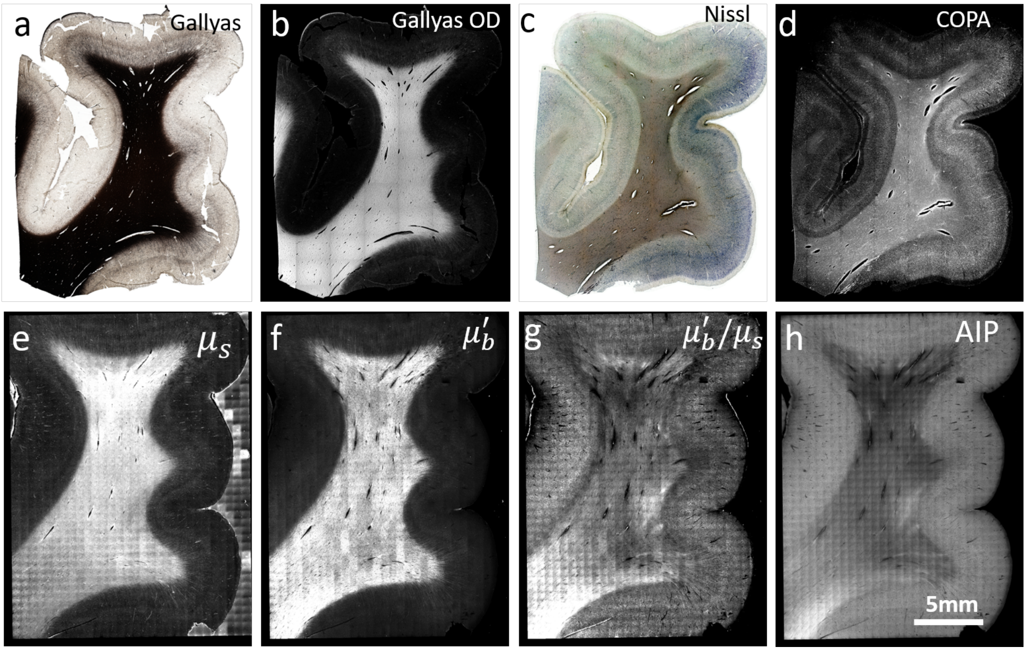


Supplementary Figure 3. Histology and optical properties of SupFrontal. (a) Gallyas Silver stain. (b) Optical density of Gallyas silver stain reveals the myelin content. (C) Nissl stain. (d) COPA. (e-g) Optical properties derived from the OCT images. (e) $\mu_s$ map. (f) $\mu'_b$ map. (g) ratio map of $\mu'_b/\ \mu_s$. (h) AIP image

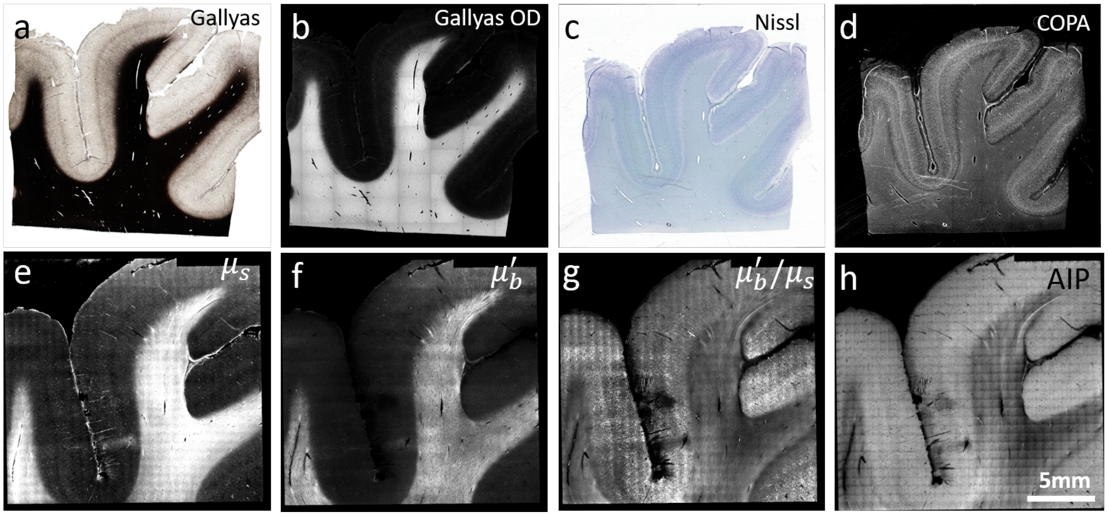


Supplementary Figure 4. Histology and optical properties of BA21. (a) Gallyas Silver stain. (b) Optical density of Gallyas silver stain reveals the myelin content. (C) Nissl stain. (d) COPA. (e-g) Optical properties derived from the OCT images. (e) $\mu_s$ map. (f) $\mu'_b$ map. (g) ratio map of $\mu'_b/\ \mu_s$. (h) AIP image

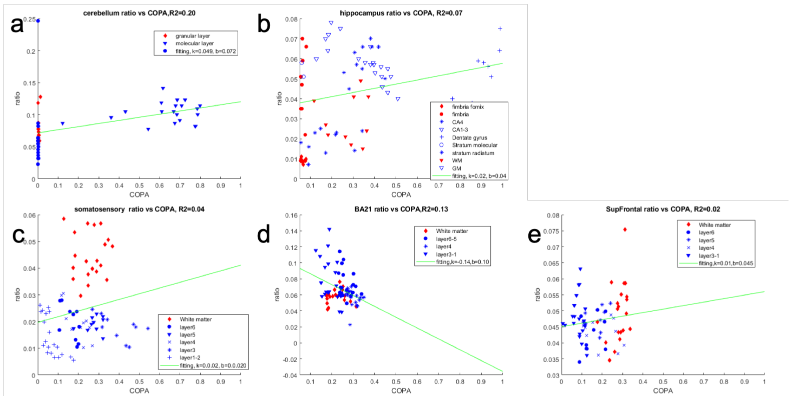


Supplementary Figure 5. Ratio of $\mu'_b/\mu_s$ of 5 samples with regard to COPA. No clear correlation between the ratio and COPA was found. Red: white matter. Blue: grey matter. Green: linear regression. (a) cerebellum. (b) hippocampus. (c) somatosensory. (d) middle temporal area 21. (e) Superior frontal

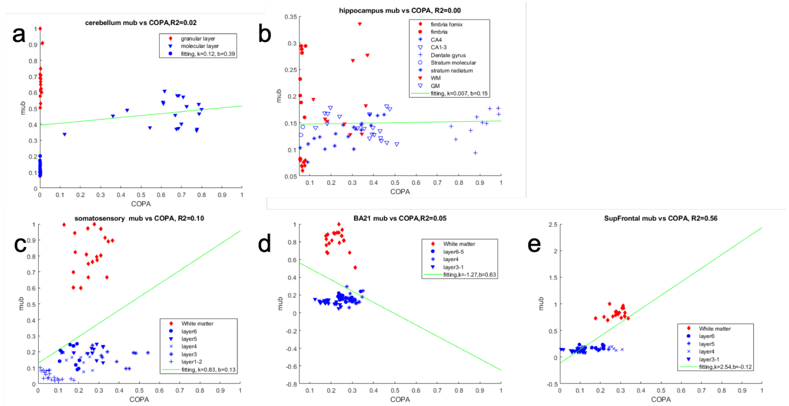

Supplementary Figure 6. $\mu'_b$ of 5 samples with regard to COPA. No clear correlation between the mub and COPA was found. Blue: grey matter. Green: linear regression. (a) cerebellum. (b) hippocampus. (c) somatosensory. (d) middle temporal area 21. (e) Superior frontal

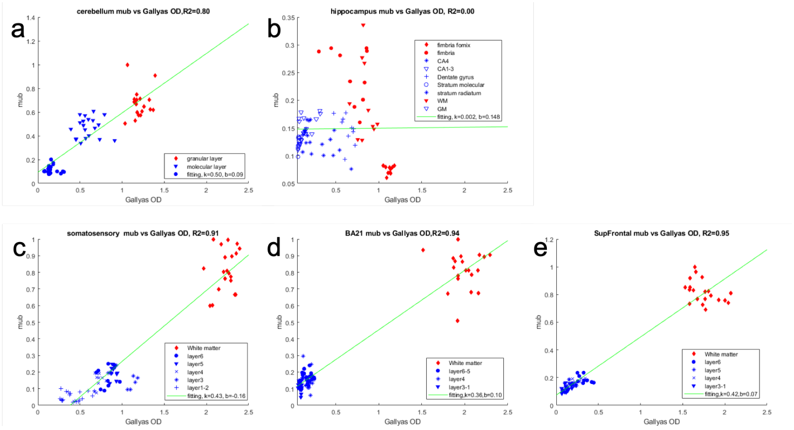

Supplementary Figure 7. $\mu'_b$ of 5 samples with regard to Gallyas OD. The cerebellum, somatosensory, middle temporal area 21 and Superior Frontal has a nice linear relationship between mub and Gallyas OD. Red: white matter. Blue: grey matter. Green: linear regression. (a) cerebellum. (b) hippocampus. (c) somatosensory. (d) middle temporal area 21. (e) Superior frontal

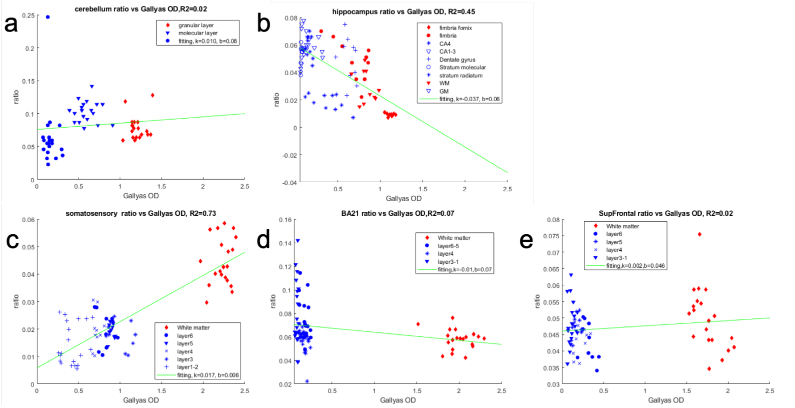

Supplementary Figure 8. Ratio of $\mu'_b/\mu_s$ of 5 samples with regard to Gallyas OD. No clear correlation between the ratio and Gallyas OD was found. Red: white matter. Blue: grey matter. Green: linear regression. (a) cerebellum. (b) hippocampus. (c) somatosensory. (d) middle temporal area 21. (e) Superior frontal

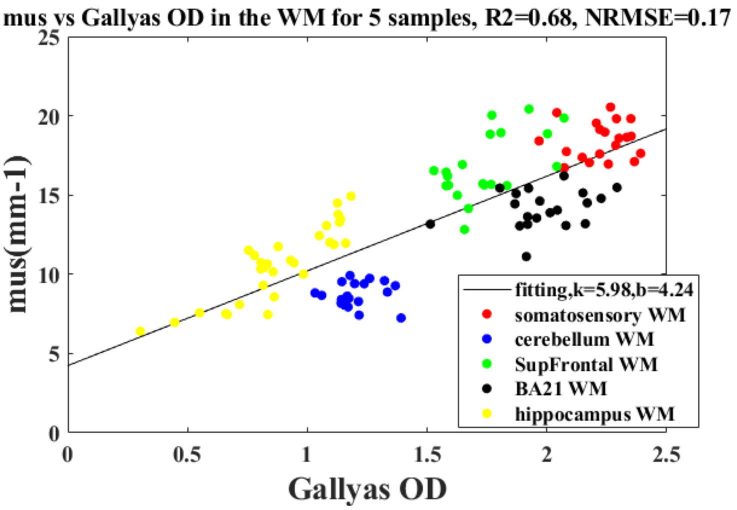

Supplementary Figure 9. 1D fitting of the white matter mus from all five samples to the Gallyas OD. Fitting shows a slope of about 6, within the slope range in Figure 2
